# Extreme copy number variation at a tRNA ligase affecting phenology and fitness in yellow monkeyflowers

**DOI:** 10.1101/392183

**Authors:** Thom Nelson, Patrick Monnahan, Mariah McIntosh, Findley R. Finseth, Kayli Anderson, Evan MacArthur-Waltz, John K. Kelly, Lila Fishman

## Abstract

Copy number variation (CNV) is a major part of the genetic diversity segregating within populations, but remains poorly understood relative to single nucleotide variation. Here, we report on a tRNA ligase gene (Migut.N02091; RLG1a) exhibiting unprecedented, and fitness-relevant, CNV within an annual population of the yellow monkeyflower *Mimulus guttatus*. RLG1a variation was associated with multiple traits in pooled population sequencing (PoolSeq) scans of phenotypic and phenological cohorts. Resequencing of inbred lines revealed intermediate frequency three-copy variants of RLG1a (*trip+;* 5/35 = 14%), and *trip+* lines exhibited elevated RLG1a expression under multiple conditions. *trip+* carriers, in addition to being over-represented in late-flowering and large-flowered PoolSeq populations, flowered later under stressful conditions in a greenhouse experiment (P < 0.05). In wild population samples, we discovered an additional rare RLG1a variant (*high+)* that carries 250-300 copies of RLG1a totaling ∼5.7Mb (20-40% of a chromosome). In the progeny of a *high+* carrier, Mendelian segregation of diagnostic alleles and qPCR-based copy counts indicate that *high+* is a single tandem array unlinked from the single copy RLG1a locus. In the wild, *high+* carriers had highest fitness in two particularly dry and/or hot years (2015 and 2017; both p < 0.01), while single copy individuals were twice as fecund as either CNV type in a lush year (2016: p < 0.005). Our results demonstrate fluctuating selection on CNVs affecting phenological traits in a wild population, suggest that plant tRNA ligases mediate stress-responsive life-history traits, and introduce a novel system for investigating the molecular mechanisms of gene amplification.

## INTRODUCTION

The standing genetic variation for fitness-related traits is a key determinant of a population’s evolutionary potential, providing the raw material for adaptive evolution when environments change (Barrett & Schluter 2008; Hoffmann & Sgrò 2011; Anderson 2016). However, the abundant genetically-based fitness polymorphism seen within many populations is also paradoxical: at any given locus, efficient directional or stabilizing selection should winnow out all but the variant with highest geometric mean fitness. Balancing selection, in which context-dependence effects of alleles on fitness maintain polymorphism, resolves this problem. Long-term protected polymorphism is robustly predicted under models of heterozygote advantage (overdominance) (Gillespie 1984) and frequency-dependent selection (Wright 1939; Clarke 1979), as well sexual antagonism (Connallon & Clark 2014) and gamete-zygote conflict (Immler *et al.* 2011; Fishman & Kelly 2015). Empirical examples of such protected polymorphism, often involving multi-gene structural variants with suppressed recombination in heterozygotes, are increasingly common (Joron *et al.* 2011; Fishman & Kelly 2015; Barson *et al.* 2015; Llaurens *et al.* 2017). However, these appealing intrinsic mechanisms of balancing selection likely apply to only a small subset of traits and genes. Spatial and (especially) temporal shifts in selection are more controversial as sources of intra-population polymorphism, but their ubiquity creates abundant opportunities to contribute to the maintenance of functional genetic variation within natural populations (reviewed in Bell 2010; Delph & Kelly 2013). Furthermore, new models that incorporate biologically-realistic factors that buffer against allele loss, such as switches in dominance (Posavi *et al.* 2014; Wittmann *et al.* 2017), plasticity (Gulisija *et al.* 2016) and the storage effect of unselected life-stages (Ellner & Hairston 1994; Svardal *et al.* 2015), suggest that environmental variation can maintain a large pool of polymorphism. However, although a few recent studies demonstrate variable selection on intermediate-frequency genetic variants (Kerwin *et al.* 2015; Chakraborty & Fry 2016; Lee *et al.* 2016; Wittmann *et al.* 2017), more work is needed to reveal whether and how environmental fluctuations contribute to functional polymorphism within natural populations.

Genic copy number variants (CNVs), including duplication and deletion of entire genes (Hastings *et al.* 2009), are likely to be functionally important components of standing variation across diverse taxa (Żmieńko *et al.* 2014; Marroni *et al.* 2014; Chain *et al.* 2014; Zarrei *et al.* 2015; Salojärvi *et al.* 2017). Changes in copy number are a common form of mutation — in plants, segmental or tandem duplication events occur at a per-gene rate at least as high at the per-site nucleotide mutation rate (Lynch & Conery 2000). Much of this variation appears to persist in populations; for example, diverse maize lines differ in copy number at nearly 35% of genes (Chia *et al.* 2012) and >15% of Arabidopsis genes are stabilized tandem duplicates (reviewed in Żmieńko *et al.* 2014). Initially, whole gene duplications or deletions may directly alter expression (Stranger *et al.* 2007), though various forms of dosage compensation can buffer these immediate effects (Veitia *et al.* 2013). Duplicates can develop novel or subdivided functions over time (Lynch & Conery 2000), but they may also influence phenotypic variation within and among populations as CNVs on shorter timescales. In plants, CNVs contribute to rapid evolution under experimental thermal stress (DeBolt 2010), underlie the contemporary evolution of glysophate-resistance in weeds (Gaines *et al.* 2010), and have been implicated in phenological and edaphic (soil stress) adaptation across populations (Rosloski *et al.* 2010; Maron *et al.* 2013; Hanikenne *et al.* 2013; Gordon *et al.* 2017). Over the longer term, CNV caused by reciprocal deletion of non-tandem duplicates is a common source of epistatic Dobzhansky-Muller incompatibilities (Bomblies *et al.* 2007; reviewed in Fishman & Sweigart 2018). However, because CNVs are difficult to assay with reference-based short-read resequencing approaches (Hunter *et al.* 2013; Tiffin & Ross-Ibarra 2014), we are only beginning to understand their contribution to within-population variation, local adaptation, and speciation. Here, we identify and characterize copy number variants of a gene contributing to the abundant phenological and fitness variation within the Iron Mountain (IM) population (Central Cascades, OR, USA) of yellow monkeyflower (Mimulus guttatus, Phrymaceae). This high-elevation annual population exhibits extremely high levels of sequence and structural diversity (Puzey *et al.* 2017) and quantitative genetic variation in fitness-relevant traits (Kelly & Willis 2001; Scoville *et al.* 2009; Bodbyl Roels & Kelly 2011; Scoville *et al.* 2011); the latter appears due in large part to intermediate frequency polymorphism, as expected under local balancing selection rather than mutation-selection or migration-selection balance against unconditionally deleterious variants (Kelly & Willis 2001; Kelly *et al.* 2013). Microsite and climatic variation (and their interaction) are strong candidates as mechanisms of balancing selection at IM, as the population is dependent on ephemeral, and highly variable, winter snowpack for summer growth and reproduction. Previously, a field study of flower-size-QTL introgression lines demonstrated fluctuating selection on two loci that affected flower size via developmental rate: in wetter years, large-flowered, relatively slowly-developing genotypes had higher fitness, but in dry years they had lower survival (Mojica & Kelly 2010; Mojica *et al.* 2012). Such antagonistic pleiotropy across environments is key to the models for the selective maintenance of standing variation in space (i.e., local adaptation in the face of gene flow) and time (Wittmann *et al.* 2017; Brown & Kelly 2018). More recently, we used PoolSeq to identify SNPs and chromosomal structural variants associated with flower size evolution under artificial selection (Kelly *et al.* 2013) and with flowering time in the field in two populations (Monnahan *et al.* 2015; Monnahan & Kelly 2017). These studies point to climatic fluctuations as a likely factor in the maintenance of standing variation within annual *M. guttatus*, but captured only a subset of the life-history traits potentially under selection.

In this study, we conducted CNV-focused reanalyses of existing population sequencing (PoolSeq) datasets for flower size (Kelly *et al.* 2013) and flowering time (Monnahan & Kelly 2017) and bulked segregant experiments (also using PoolSeq) for vernalization requirement and germination time in the field. Both of these traits may mediate the timing of reproduction (a key trait for plant fitness), and also affect the probability of survival to flowering under winter-and drought-stress. The timing of germination may be a strong determinant of the fitness effects of genes underlying flower size and development speed. Because multiple population contrasts pointed to copy number variation at a tRNA ligase gene (Migut.N02091, RLG1a) as a correlate of life history and phenological variation, we then characterize RLG1a CNVs using resequencing of inbred lines, qPCR of genomic DNA, and PCR-based genotyping. One of these variants (found at intermediate frequency in inbred lines and field samples) has two additional copies of RLG1a (*trip+),* while the other (absent from a large inbred line set, and at low frequency in the wild), has ∼300 ampliconic copies (*high+).* Using markers diagnostic of these two multi-copy genotypes, we demonstrate the expected associations between individual CNV genotypes and phenology/flower size in multiple PoolSeq experiments. In independent experiments, we show that carriers of the *trip+* variant exhibit elevated RLG1a expression and that they delay flowering under experimentally water-stressed conditions. Finally, we examine the frequency and fitness effects of the three CNV genotypic categories over three years spanning extremes of fecundity and survival in the Iron Mountain population, finding significant (but variable) associations between genotype and fitness in each year. Together, these analyses suggest that copy number variants of a *Mimulus guttatus* tRNA ligase homolog influence life history traits that experience fluctuating climatic selection.

## METHODS

### Study System

The *Mimulus guttatus* (also known as *Erythranthe guttata*; Phrymaceae) species complex exhibits tremendous diversity in life history, mating system, and other traits across Western North America. *Mimulus guttatus*, the most widespread and diverse species (and the likely progenitor of other complex members) is self-compatible but largely outcrossing (Sweigart *et al.* 1999). We focus here on the well-studied Iron Mountain (IM, Oregon) high-elevation annual population, from which the inbred line (IM62) defining the *M. guttatus* reference genome was derived. This population exhibits high levels of standing genetic variation for fitness-related traits (including flowering time and flower size) (Kelly *et al.* 2013; Monnahan & Kelly 2017) as well as multiple segregating inversions with opposing effects on fitness (Fishman & Kelly 2015; Lee *et al.* 2016). These features reflect a very large population (census size in 100,000s each year) without internal genetic structure (Sweigart *et al.* 1999) and dominated by natural selection rather than drift (Puzey *et al.* 2017). Ecologically, the life cycle of the IM population is driven by variable yearly patterns of snow accumulation and melt-out, as these determine the timing, length, and lushness of the summer growing season. Germination occurs in both Fall (November) and in spring after melt-out (May-June, generally). Plants die by drying out, and have a growing season of five to twelve weeks depending on shared interannual variation in snowpack and weather, as well as individual microsite characteristics (e.g., soil depth). IM and nearby sites exhibit high variance in mean fitness (both survival and fecundity components) from year to year (Mojica *et al.* 2012; Fishman & Kelly 2015; Monnahan *et al.* 2015). Phenotypic selection analysis (Fishman & Willis 2008) and field experiments with introgression lines (Mojica & Kelly 2010; Mojica *et al.* 2012) indicate that plants with large flowers are strongly favored, assuming they live to flower. However, negative correlations between flower size and development rate (i.e., big = slow) make genetically large-flowered plants more likely to die before flowering in dry years (Mojica & Kelly 2010; Mojica *et al.* 2012). This sets the stage for fluctuating selection on loci that mediate or modulate this tradeoff (Monnahan & Kelly 2017; Brown & Kelly 2018).

### PoolSeq of phenotypic populations

To screen for copy-number variants contributing to life history variation genome-wide, as well as fully characterize the focal RLG1a locus, we re-analyzed PoolSeq datasets from two previous experiments. The High-Low-Ancestral set (Kelly *et al.* 2013) included large-and small-flowered populations derived by nine generations of artificial selection with outbreeding (High and Low selection lines; pool N = 49 and 78, respectively), starting from a large outbred IM-derived base population (Ancestral, pool N = 75). A Control population maintained under the same conditions, but without selection, was also generated (Kelly *et al.* 2013); it was not sequenced, but we use it here for individual genotyping tests. The Early-Late set consisted of field-collected cohorts of early-and late-flowering plants from IM and nearby Quarry populations, from both 2013 and 2014 (Monnahan & Kelly 2017). Because the Quarry population is known to be the product of recent admixture between IM-like annuals and perennial genotypes (Monnahan *et al.* 2015), we restrict our analyses here to the IM datasets.

Following Monnahan & Kelly (2017), we constructed two new pairs of (PoolSeq) libraries, each contrasting alternative phenological/trait “populations” within the outbred Iron Mountain population. *Fall-Spring*: In fall 2012 and spring 2013, we collected fall-germinating and spring-germinating (cotyledons only) seedlings from nine quadrats spaced across the IM site (N= 40 samples per quadrat collected 20/tube, except for quadrant in Spring with only 10 seedlings; total N = 320 and 290, respectively.) After extraction, tubes were combined proportionately (equimolar concentrations by sample size) into Fall and Spring pools for library preparation. *Bolt-Rosette.* In spring 2012, we grew >600 independent (1 per maternal family) plants derived from 2011 field-collected seeds in University of Montana greenhouse under non-inductive daylengths (12 hour days) for 2 months, and then transitioned to 16 hour days. These conditions induce flowering in IM *M. guttatus* that do not require vernalization (a cold treatment simulating winter) and inhibit it in those that do (Friedman & Willis 2013; Fishman *et al.* 2014). About 60% of plants had flowered after 60 days at 16hr daylengths. Individuals were then assigned to Bolt (flowered without vernalization) and Rosette (vernalization-requiring, non-flowering) cohorts, and tissue collected in sets of 20 plants per extraction tube. After DNA extraction using a standard Mimulus protocol (Fishman & Kelly 2015), 10 and 15 tubes per population were pooled in equimolar amounts (N = 300 and 200 plants per Bolt and Rosette pools, respectively).

### PoolSeq and Line Resequencing and Analysis

The High-Low-Ancestral flower size and Early-Late flowering time cohorts were sequenced as described previously (Kelly *et al.* 2013; Monnahan & Kelly 2017). The Fall-Spring and Bolt-Rosette population pairs were each sequenced with Illumina HiSeq 2500 (PE 100) at the University of Kansas Genomics Core, following the same library preparation protocol as previous pools (Monnahan & Kelly 2017). All eleven PoolSeq populations, and reanalyzed IM inbred lines, were processed with the same bioinformatics pipeline (see Supporting Information for details).

For analyses of coverage differences among pooled populations and lines, we restricted our analyses to sites in exons, as extremely high diversity within *M. guttatus* hampers read-alignment outside of genes (Puzey *et al.* 2017). We calculated mean read depth at each annotated exon with Mosdepth (Pedersen & Quinlan 2018) and then standardized exon read-depth values by the genome-wide median for a given pool or sequenced individual. After excluding a subset of annotated genes that appeared to be chloroplast-nuclear transfers (or mis-assembled), and a set that appeared to be mis-annotated repetitive DNA (see supplemental methods in Supporting Information), we retained 25019 genes with coverage data from one or more exons. To compare read depths among pools genome-wide, we calculated the absolute value of the difference in standardized read depth between each pair of populations (e.g., High vs. Ancestral, Low vs. Ancestral, Bolt vs. Rosette, Spring vs. Fall, IM13_Early vs. IM13_Late, IM14_Early vs. IM14_Late). For comparisons of mean RLG1a coverage among pools, we restricted estimates to the final 23 exons (spanning ∼9 kilobases) of Migut.N02091, as the first three exons showed high variance in coverage depending on whether or not unpaired reads were filtered out and we cannot definitively distinguish additional duplication of this portion of the gene from mis-mapping.

Using a similar bioinformatics pipeline, we examined Migut.N02091 exon coverage in previously resequenced IM inbred lines (1 from Lee *et al.* 2016; 34 from Puzey *et al.* 2017) and the outbred *high+* lines (see below). To identify sequence variation among genomic copies in trip+ individuals, we performed metagenome *de novo* assembly on IM664 sequence reads mapping to Migut.N02091. After converting the SAM alignments for extracted reads to fastq format, we used the metagenome assembler MEGAHIT (Li *et al.* 2015) with default settings. Assembled coting’s were mapped back to the *Mimulus guttatus* version 2 reference genome with minimap2 (Li 2018)to identify haplotypes diagnostic of duplicated copies of the gene.

### Genetic marker design and genotyping

For analyses of individual genotype, we designed and amplified an exon-primed length-polymorphic marker (mN2091×16) spanning a 5 amino-acid (5AA; 15 base) insertion-deletion polymorphism in Exon 16 of Migut.N02091, as well as the highly variable Exon16-17 intron (Supporting Information: Table S1). In inbred lines (N = 14 tested) and wild-derived plants, we found multiple (diploid) amplicons corresponding to single-copy sequences (RLG1a_*solo*), plus two amplicons carrying the Exon 16 5AA deletion: a 426bp fragment was exclusive to IM lines with ∼3x coverage of Migut.N02091 (*trip+)* and a rare 428bp amplicon was uniquely associated with very high coverage individuals (*high+)* found in wild collections. Both 5AA-deletion variants at mN2091×16 were scored as presence/absence polymorphisms, as the *high+* 428bp allele was always “homozygous” due to out-competition of all other alleles in PCR reactions and the *trip+* 426bp allele was always found with 2-3 additional amplicons also found in *solo* lines (see Results).

### Characterization of RLG1a trip+: vernalization, RLG1a expression, and flowering time under stress

To test whether the patterns observed the PoolSeq results from shifts in RLG1a CNV frequencies, we genotyped individual plants from the flower size selection (High, Low, and Control) populations and from IM14 (Early and Late flowering) populations at mN2091×16 marker. We also performed two separate experiments to assess phenological phenotypes. We grew *solo* and *trip+* inbred lines (inferred from marker genotyping) under the same greenhouse regime as in the Bolt-Rosette PoolSeq experiment in Spring 2015 and sampled rosette leaf tissue one day prior to and six days after the transition from 12hr daylength to16hr daylengths (n = 9 *solo* lines and 2 *trip+* lines, respectively; 2-4 separate individuals per treatment per line; total N = 56). Both *trip+* lines (IM664 and IM115) flowered within one month under this regime, consistent with a lack of vernalization requirement, as did three of the 10 *solo* lines. We extracted RNA, made cDNA, and measured expression of RLG1a (mN2091q primers in Exon 26) relative to two previously developed controls (EF1a and UBQ5) (see Supporting Information for details). Differences in expression were determined from threshold cycle number (Ct) relative to the control (ΔCt = Ct _mN2091_ - Ct _EF1a;_ averaged over 2 technical reps each) and analyzed using ANOVA with daylength (12hr, 16hr), RLG1a CNV category (*solo, trip+*), and line (nested within CNV) as main effects and the and daylength x CNV interaction effect.

As part of a separate study on differentiation in soil-microbial interactions, we grew IM outbred plants (from 2013 wild seed) in the University of Montana greenhouse in Summer 2017 in two soil treatments: soil from the Oregon Dunes perennial *M. guttatus* site (DUN) (Hall & Willis 2006) and from IM, each cut 50:50 with sand. Plants were top-watered daily and fertilized weekly with a low-phosphorus fertilizer. Both treatments mimicked drought-stressed field conditions and produced rapid-flowering phenotypes resembling wild IM plants. We infer the DUN soil to be more stressful because it produced a ∼2-fold reduction in mean biomass and fruit number at harvest vs. the IM soil (data not shown). We recorded the number of days to first flower, extracted DNA from IM plants, and genotyped individuals at the mN2091×16 marker. Flowering time was analyzed with ANOVA with soil type and genotype (*trip+* /*solo*; three *high+* carriers were excluded from this analysis) and their interaction.

### Characterization of RLG1a high+ variants: resequencing, qPCR, and genetic segregation

To characterize rare high copy (*high+)* variants of RLG1a, we grew and PCR-genotyped plants from 2013 and 2015 seed collections (IM13P, IM15P; N = 120 maternal families total, 1-4 progeny each). One *high+* individual from each year (IM13P_627c and IM15P_317d, respectively) was chosen for whole genome sequencing. Library preparation, sequencing (HiSeq 4000, PE125), and alignment to the *M. guttatus* V2 reference are described in Supporting Information. Median-standardized exon coverage for Migut.N02091 was calculated as with the PoolSeq and line datasets.

In addition, we counted RLG1a genomic copy number by qPCR in two wild-collected *high+* plants, IM13P_644 and IM13P_627c, as well as *trip+* and *solo* lines. As single-copy control, we used primers within Exon 12 of Migut.G00571 (Isopropyl Malate Isomerase large subunit; mG571q in Table S1). RLG1a ΔCt values (Ct _[mN209_– _Ct[mG571]_], were calculated for each sample, averaged cross two technical replicates, and copy number estimated using the equation: 2^(Δ Ct solo - Δ Ct high+)^. To corroborate these estimates in IM13P_644 (which had highest copy number), we created a standard curve by performing 3-fold serial dilutions of template DNA and measuring Ct for each dilution. We then used the linear fit of the standard curve to estimate mN2091q copy number relative to mG571q in the same plant.

To segregate the high copy allele, we selfed IM13P_627c and grew the resultant progeny (which segregate as a pseudo-F_2_ at loci heterozygous in the parent plant) in the University of Montana greenhouse in Spring 2018. All progeny (N = 192) were genotyped with mN2091×16 to score *high+* presence/absence. For tests of location, we also genotyped a linkage-informative (heterozygous in IM13P_627c) marker <100kb from Migut.N02091 (mN2089; Table S1) on a subset. To confirm segregation of the *high+* cluster as a single locus, we conducted genomic qPCR (as above) on a subset of progeny (N = 39). If all copies are in tandem, *high+* carriers should fall into two distinct categories with a ∼2-fold difference in copy number. Individual ΔCt values (calculated as above) were clustered using Normal Mixture methods, the cluster number with the lowest AIC/BIC value chosen, and cluster means compared with ANOVA. Finally, to assess whether expression was elevated in high-copy carriers, we also conducted rt-qPCR on mRNA extracted from floral shoot tissue of a small set of progeny (segregating for 0, 1 or 2 *high+* alleles based on the genomic DNA qPCR), using EF1a as a control as in the *trip+* analyses. Additional methods in Supporting Information.

### Frequency and fitness in the wild

To evaluate RLG1a CNVs in the field, we genotyped and measured fitness of wild IM plants in 2015-2017. In 2015, we marked spring (cotyledons) and fall (rosette leaves) germinants in May, but almost all of the spring germinants died as seedlings and were lost before DNA could be collected. We marked a second set of flowering plants on two dates a week apart in mid-June; most of these died as well, but we obtained DNA from many non-survivors. Finally, we collected tissue for DNA extraction and all seeds from survivors among the marked germinants and flowerers, as well as additional random survivors as in previous years (Fishman & Kelly 2015). Thus, we can estimate both survival from flowering to fruit and seedset among survivors, from which we can calculate lifetime fitness conditioned on surviving long enough to be collected. In 2016 and 2017, we obtained DNA and fruit/seed counts of plants that survived to fruit, as in previous studies. Individuals were genotyped at mN2091×16, and coded as *trip+, high+* or *solo*. We analyzed log-transformed lifetime female fitness (survival x seedset +1) from 2015 alone using standard least squares ANOVA. We analyzed seedset and fruitset from 2016 and 2017 together, with GLM analyzes (Poisson, log link) including year, RLG1a genotype, and their interaction.

All statistical analyses of qPCR, genotypic, and phenotypic data were performed in JMP 13 (SAS Institute 1994).

## RESULTS

### PoolSeq reveals RLG1a coverage variation associated with multiple life history traits

A tRNA ligase gene, Migut.N02091 (RLG1a), was a strong read-depth outlier in four of the six PoolSeq comparisons (Fig. 1), including the IM14_Early vs. IM14_Late comparison in which it was previously identified as a SNP outlier (Monnahan & Kelly 2017). The 11 PoolSeq populations varied widely in genome-standardized coverage across Migut.N02091 (Exons 6-28), ranging from near-background in the Ancestral (mean = 1.18) to >10x in the Bolt and Spring germinant pools (means = 11.20 and 11.07, respectively). Like the Ancestral population for flower size selection, both IM13 flowering cohorts, IM14_Late, and the Low flower size selection population exhibited relatively low N2091 coverage (1.2-1.67x background) indicating few individuals with high-copy genotypes in these pools. The High flower size selection population, Fall germinant, IM14_Early, and Rosette pools were intermediate in coverage (5.82, 5.99, 6.70, and 3.1, respectively).

**Fig. 1.**
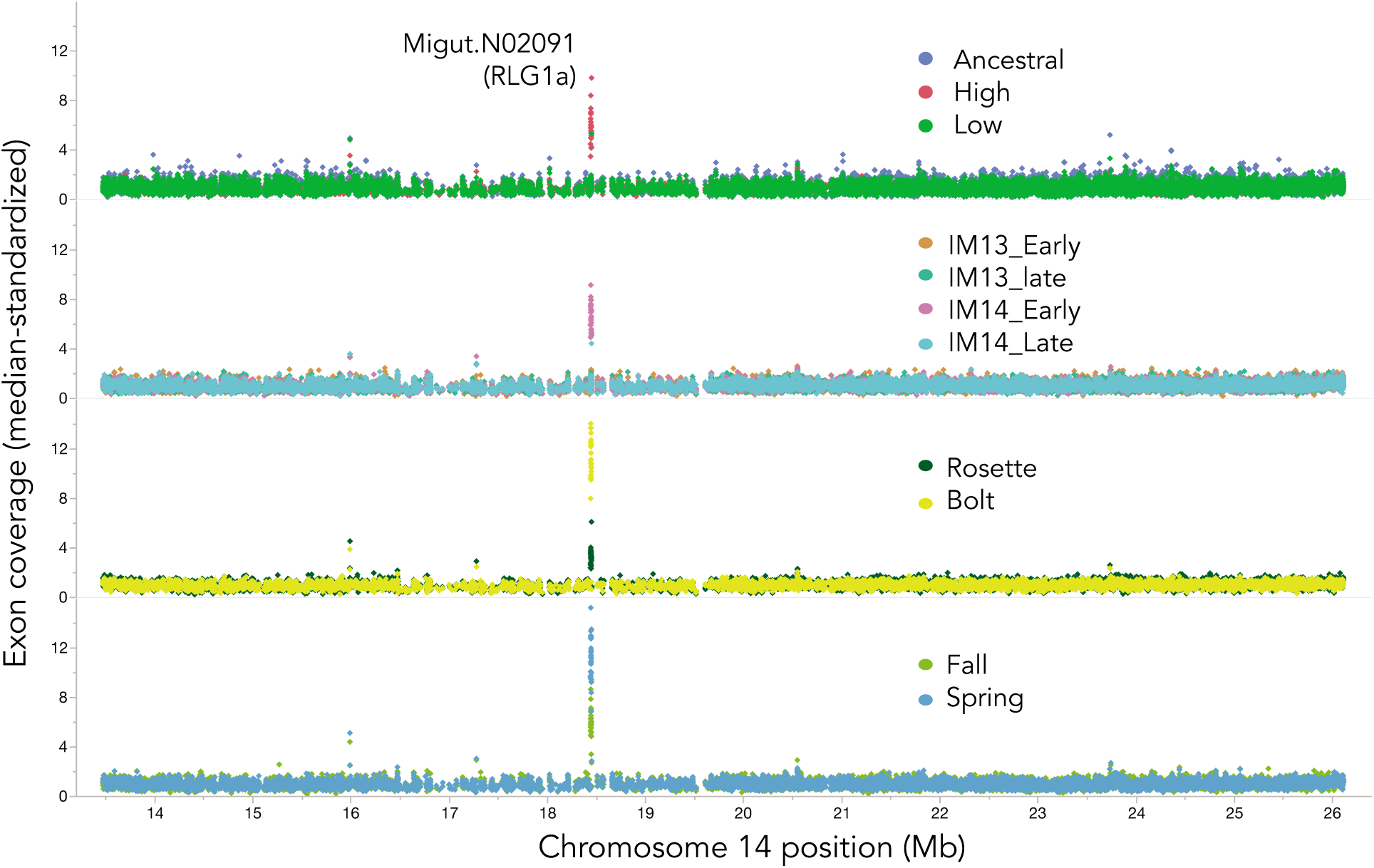
Median-standardized exon coverage (read depth) in pooled resequencing of 11 Mimulus guttatus phenotypic populations, shown for 13Mb of Chromosome 14 flanking focal gene RLG1a.

### A common 3-copy (trip+) variant of RLG1a contributes to flower size and flowering time variation

Differences in standardized coverage of RLG1a between paired pools (Fig. 1) suggest shifts in the frequency of multi-copy haplotypes between cohorts/phenotypes. To identify haplotypes contributing to these PoolSeq patterns, we examined sequence variation and RLG1a exon coverage in a set of 35 previously re-sequenced inbred lines from IM. Four lines (IM664, IM115, IM922, IM239) exhibited a characteristic pattern of ∼3x elevated coverage (and heterozygosity) across most of the gene (last 23 exons) plus extremely high coverage (26-47x background) of the first three exons (Fig. 2). These lines were uniquely scored as heterozygous for a 15bp (5 amino acid; 5AA) deletion in Exon 16 relative to the reference sequence plus three linked intronic SNPs. Metagenomic assembly of IM664 identified three haplotypes across this section of the gene; one of these contains the 5AA Exon 16 deletion and linked SNPs. One additional line (IM1145) exhibited a similar (but weaker) pattern of elevated coverage gene-wide (mean = 16.4x across first 3 exons and remainder of gene, respectively) and was only heterozygous at the SNPs. We infer it to be a heterozygote of multi-and single-copy genotypes at the time of resequencing. Thus, 1/7 of the inbred lines carry an RLG1a genotype (henceforth *trip+*) with three distinct paralogs plus (apparently) additional duplications of the 5′ end of the gene. All *trip+* lines tested (n = 4, including IM1145) carried a 426bp mN2091×16 amplicon (with 5AA Exon 16 deletion), which as absent from single-copy *(solo)* lines (n = 9). *trip+* lines (which are highly inbred and should be homozygous genome-wide) always had at least two other amplicons, which differed among *trip+* plants and overlapped in size with single-copy (*solo)* amplicons. Therefore, we score *trip+* as a dominant (presence/absence) marker. An additional four inbred lines (IM138, IM359, IM549, IM709) exhibited ∼2x coverage (Fig.2); and dozens of heterozygous sites gene-wide together, this suggests stacking of diverged duplicates rather than retained heterozygosity. Unfortunately, we were unable to identify diagnostic marker alleles for this putative RLG1a_*dup* genotype; therefore we roll them into the single-copy (*solo*) genotypic class for analyses of outbred individuals.

**Fig. 2.**
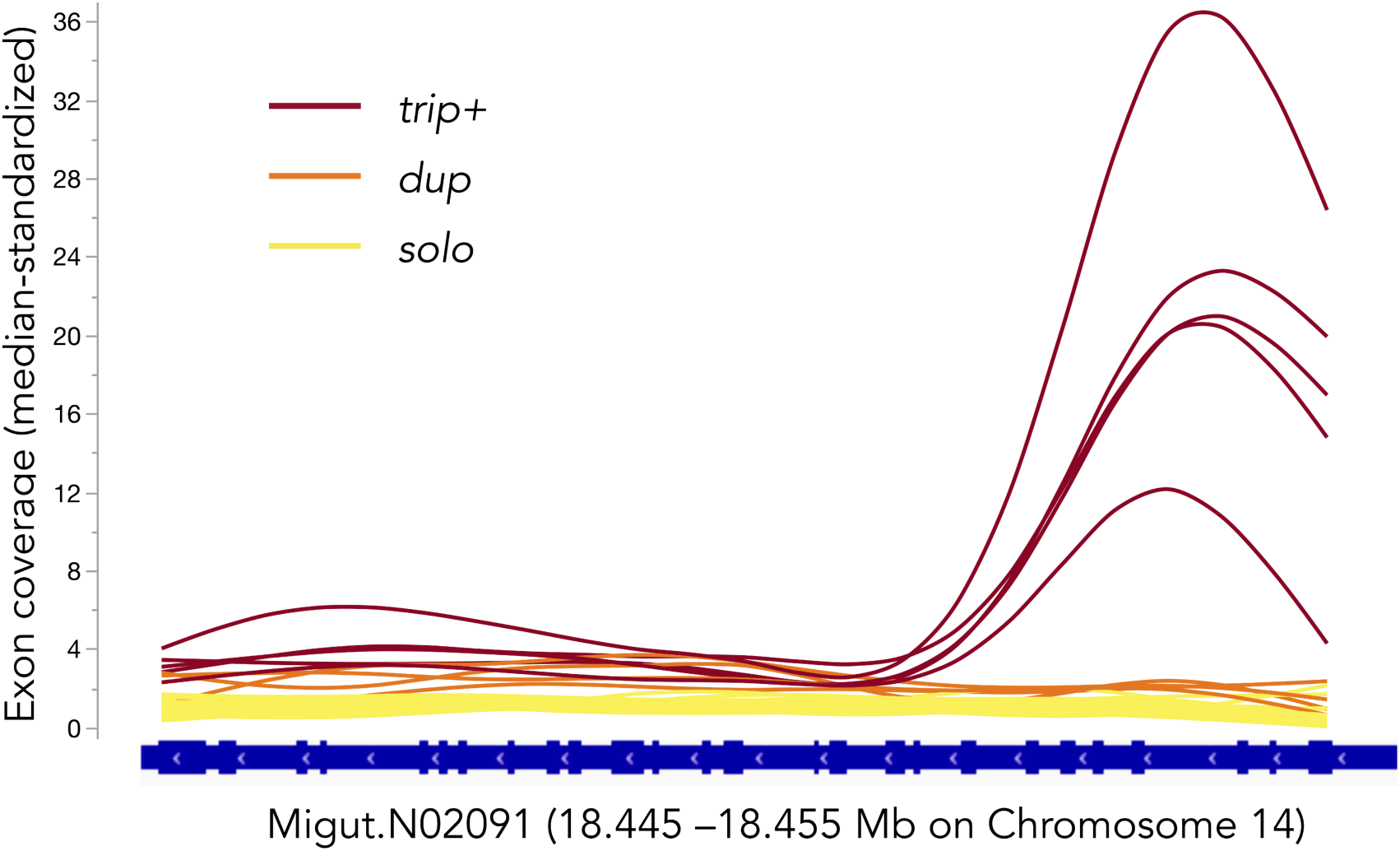
Smoothed exon coverage (standardized by chromosomal median) of Migut.N02091 (RLG1a) in 35 inbred lines of Iron Mountain *Mimulus guttatus*. Colors indicated inferred CNV category based on gene-wide coverage patterns and “heterozygosity”. The marker mN2091×16 spans between exons 16-17 and the qPCR marker mN2091q is in exon 26. The gene region containing the first three exons shows homology to the housekeeping gene EF1a, and may be additionally duplicated in *trip+* individuals.

Individual genotyping confirmed a substantial (x^2^_2 df_ = 11.2, P < 0.005) difference in *trip+* frequency among the flower size selection populations, with twice as many *trip+* plants in the High (36/72 = 50%) vs. Low (13/55 = 26%) and Control (7/29 = 24%) cohorts individually genotyped. Plants carrying *trip+* also exhibited late flowering in two independent datasets, one from the field and one under field-mimicking conditions in the greenhouse. First, individual genotyping of the 2014 field PoolSeq collections revealed twice the frequency of *trip+* plants (32%) in Late vs. Early (15%) flowering cohorts **(χ^2^_1 df_** = 6.3, *P* = 0.012, N = 179). This may reflect a relatively slow transition to flowering by *trip+* plants under the drought-stress conditions experienced in normal-to-dry years at Iron Mountain. Consistent with this inference, *trip+* was associated with later flowering in plants grown under two stressful soil conditions in the greenhouse (F _3. 75_ = 6.6, *P* = 0.03 for genotype effect). Although, the interaction with soil type was nonsignificant (*P* = 0.09), the main genotypic effect was primarily due to delayed flowering of *trip+* plants (mean = 38.1 ± 1.3 SE days, n = 17) relative to non-carriers (mean = 33.4 ± 1.3 SE days, n = 18) in the sandier and more drought-stessed DUN soil, with no difference evident in the IM soil (Fig. 3a).

**Fig. 3.**
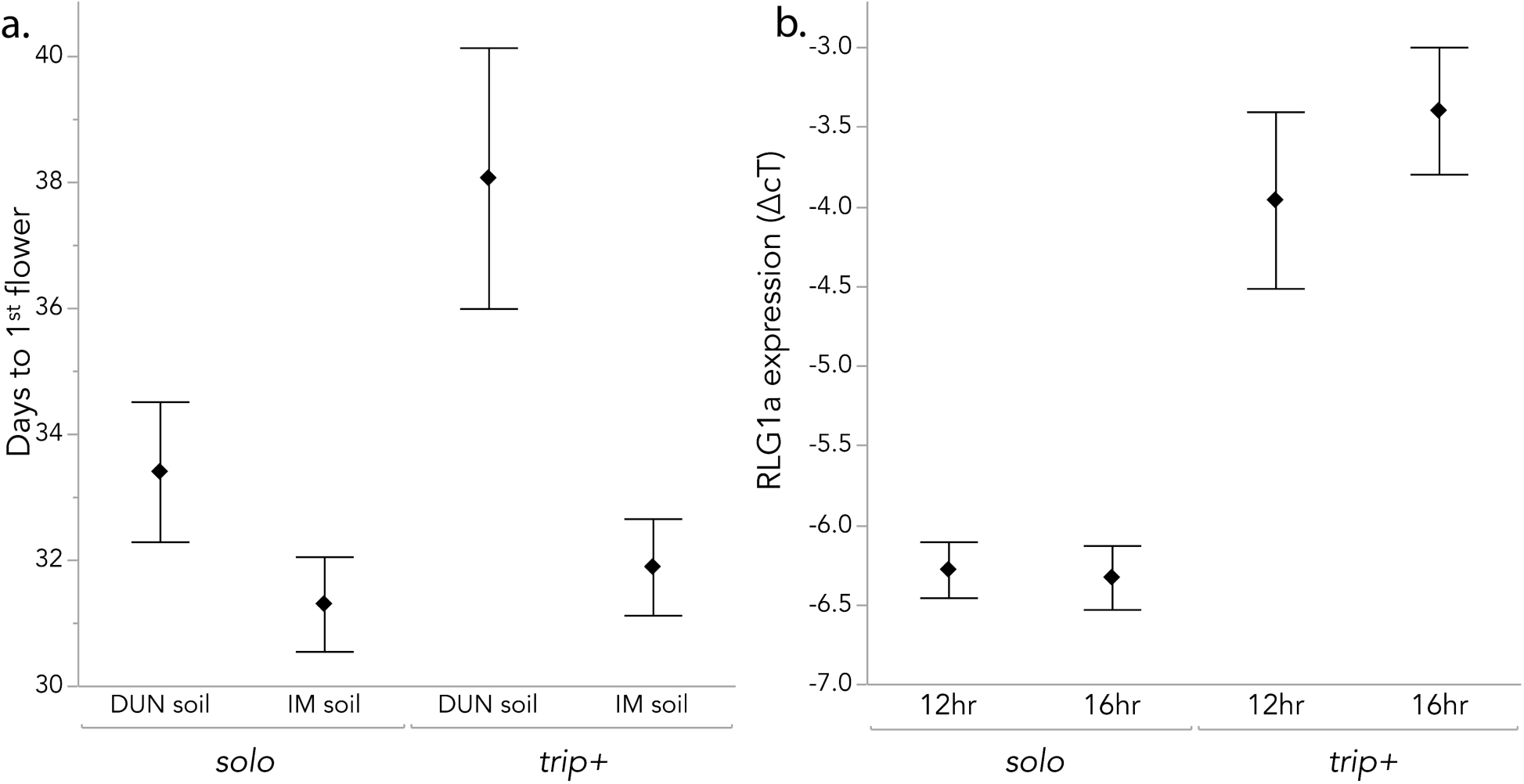
Flowering time and expression of RLG1a genotypes under controlled conditions**. a)** Time to first flower (mean ± 1 SE) for outbred plants grown in two soils in greenhouse. Analyzed with ANOVA (overall r^2^ = 0.21, n = 79). Soil type effect: P < 0.001; RLG1a genotype effect: P = 0.03; interaction: P = 0.09). **b.** RLG1a expression (relative to EF1a control; mean ± 1 SE) in leaf tissue of *solo* (N = 9) and *trip+* (N =2) inbred lines under two daylength conditions. Analyzed with nested ANOVA (overall r^2^ = 0.78, n = 56): Daylength effect: P = 0.58; RLG1a CNV effect: P < 0.0001; Line nested within genotype effect; P = 0.002)

Consistent with increased mRNA dosage due to increased copy number, two *trip+* lines exhibited highly elevated RLG1a expression relative to *solo* lines (n = 9) under both short (12hr) and long (16hr, after 12hr) daylength conditions (P < 0.00001 for RLG1 genotype effect; Fig. 3b). Daylength, non-vernalized flowering phenotype (Bolt/Rosette), and their interaction had no effect on RLG1a expression in leaves (all P > 0.15), but line was highly significant beyond the RLG1a CNV effect (P < 0.002). The RLG1a CNV effect was not a consequence of flowering phenotype; both *trip+* lines tested had the Bolt phenotype, but so did two of the *solo* lines. However, there was hint of interaction between daylength and CNV (P = 0.09), with one of the *trip+* lines in particular (IM115) exhibiting increased expression after the shift to 16hr days.

### A rare variant with ∼300 RLG1a copies (high+) drives PoolSeq coverage patterns and segregates at a single nuclear locus

Even dramatic shifts in the frequency of a 3-copy *trip+* haplotype cannot explain the >10x gene-wide coverage of RLG1a in the Spring germinant, Bolt, and IM14_Early cohorts. In genotyping individuals from the IM14 and flower size selection PoolSeq populations, we identified a second amplicon with the 5AA Exon 16 deletion (428bp: 3 each in High and Ancestral, 0 in Low; total N = 162), and in IM14_Early (6/90 = 6.7%) but not IM14_Late (0/95 = 0%). The skew in incidence of the 428bp allele in IM14 was statistically significant, even when *trip+* (elevated in IM14_Late; see above) and *solo* variants were coded as a single category (Pearson **χ**^2^ = 6.5, p = 0.01). Despite its rarity, the 428bp amplicon (hereafter *high+)* was always “homozygous”, suggesting it out-competes other alleles during mN2091×16 PCR-amplification.

None of the resequenced IM *M. guttatus* inbred lines (N = 35) had RLG1a coverage greater than *trip+* or the indel variation necessary to generate a 428bp mN2091×16 allele. However, two wild-derived 428bp carriers (IM13P_627c and IM15P_617d) exhibited a stack of ∼275x (median-standardized) coverage across Migut.N02091 (Fig. 4a). The vast majority of stacked reads represented a single haplotype, with the Exon 16 5AA deletion present at >99% (1608/1626 reads had no coverage at the central site of the deletion). Elevated coverage extended across the entire gene as well as the intergenic region between Migut.N02091 and Migut.N02090, spanning ∼19 kb. If all copies were arranged in tandem, this *high+* RLG1a haplotype would span at least 5.7Mb, or ∼22-46% of a *M. guttatus* chromosome in the current v2 assembly.

**Fig. 4.**
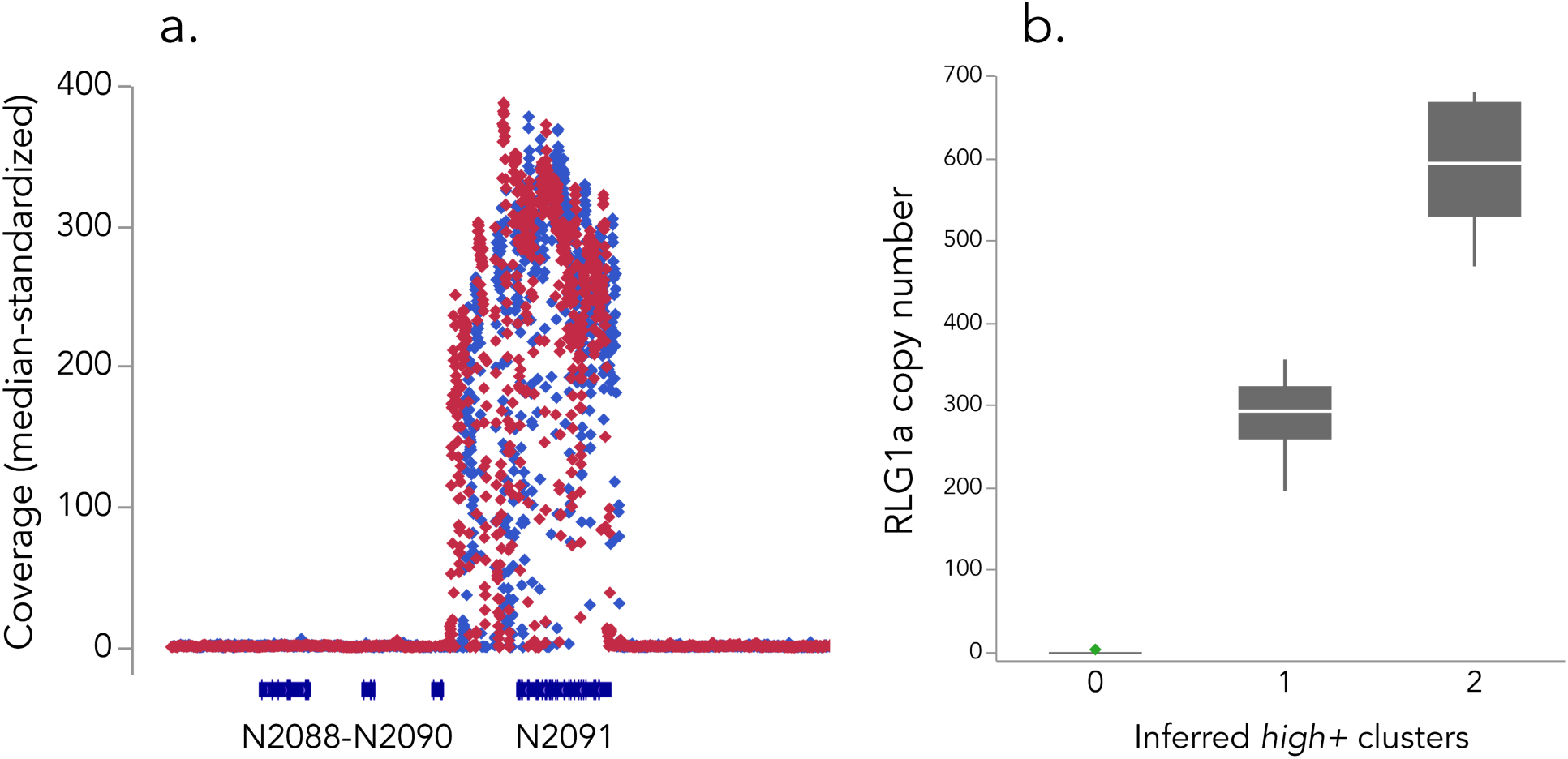
Identification and segregation of *high+* RLG1a genotype. **a.** Median-standardized coverage (25bp windows) of *high+* carriers IM13P_627c and IM15P_317d, showing amplification of Migut.N02091 and the Migut.N02190-2091 intergenic region. **b**. Quantile box plots for RLG1a copy number in IM13P_627c progeny segregating for 0 (n = 7), 1 (n = 20) or 2 (n = 12) *high+* variants. RLG1a copy number was estimated as 2(Δ Ct), where ΔCt is the absolute difference between Ct _mN2091q_ and Ct _mG571q_ for an individual, averaged across two technical replicates.

qPCR of genomic DNA confirmed a massive copy-number expansion in *high+* carriers. In two wild-derived carriers, (IM13P_627c and IM13P_644), N2091q amplified 8.5 and 9.5 cycles earlier than the single copy reference, respectively. An internally controlled standard curve estimated copy number of the latter to be >500 (502 and 513 in two technical replicates), and the single-cycle difference in ΔCt is consistent with IM13P_627c having one *high+* “allele” and IM13P_644 having two). In the PoolSeq populations, capture of fewer than 10 *high+* individuals (each with ∼250-300 copies of RLG1a), can explain elevated coverage in IM14_Early vs. IM14_Late. Although we could not genotype individuals from the Fall-Spring and Bolt-Rosette experiments, commensurate shifts in frequency (e.g., *high+* carriers relatively common in Bolt and Spring, but absent from Rosette and other low coverage pools) plausibly explain their coverage differences.

The extraordinary amplification of RLG1a in *high+* plants raises the possibility of transfer of nuclear DNA to an organellar genome (which occur in many copies per cell), rather than duplication within the nuclear genome. Furthermore, even if encoded in the nuclear genome, RLGA1 could be dispersed across many clusters genome-wide (Gaines *et al.* 2010). Finally, even if tandemly amplified as a single genetic locus, *high+* copies need not be located in tandem with (solo) reference Migut.N02091. To distinguish these possibilities from the working hypothesis that *solo* and *high+* versions of RLG1a are alternative alleles at the same locus, we examined segregation in a pseudo-F_2_ made by selfing *high+* carrier IM13P_627c. The proportion of *high+* carriers in the progeny (84% or 146/173) was slightly elevated over the Mendelian expectation of 75% for a dominant marker (**χ**^2^_2 df_ = 4.58, *P* = 0.03), but also substantially lower than the two-locus expectation of 93.75% (**χ**^2^_2 df_ = 7.57, P = 0.006). Genomic qPCR confirm the inference of a single *high+* locus: progeny ΔΔ Ct values (N = 39) clustered into three distinct groups (normal mixtures analysis, AIC/BIC lowest for 3 clusters). *Solo* individuals had ΔΔCt values not different from 0 (mean = 0.10 ± 0.14; n = 7), whereas *high+* carriers sorted into hemizygous (mean ΔΔ Ct = 8.195 ± 0.08; n = 20) and homozygous (mean ΔΔ Ct = 9.19 ± 0.11; n = 12) classes with the expected 2-fold difference in inferred copy number (286.5 ± 11.6 SE and 590.5 ±15.9 SE, respectively; Fig. 4b. Both marker and qPCR segregation ratios point to a single nuclear locus segregating 0, 1 or 2 *high+* RLG1a arrays consisting of ∼300 copies of the gene.

To test whether the *high+* locus was genetically coincident with Migut.N02091 (i.e., a local tandem expansion on Chromosome 14), we genotyped 62 IM13P_627c progeny at a tightly-linked flanking marker (mN2089). In *solo* progeny (n = 8), mN2091×16 segregated for two amplicons perfectly associated with segregating mN2089 genotype, indicating that Migut.N02091 is correctly located in the reference genome. However, mN2089 also exhibited Mendelian segregation <u>within</u> the remaining *high+* carriers (**χ**^2^_2 df_ = 1.2, *P* >0.50 vs. 1:2:1 expectation). Two segregating alleles at Migut.N2091 in *solo* progeny, plus no linkage between *high+* presence and m2089 genotype, demonstrates that RLG1a *high+* segregates as a presence/absence polymorphism at an unlinked locus. Thus, RLG1a exhibits multiple levels of structural variation, varying in the number of genetic loci as well as the number of copies per locus.

As a coarse assay of whether *high+* carriers exhibited constitutive expression of RLG1a in proportion to copy number, we conducted rt-qPCR on shoot tissue from greenhouse-grown siblings of the two sequenced *high+* plants (outbred, so segregating for all possible genotypes). Both *high+* (ΔCt = -2.87, n=3) and *trip+* (ΔCt = - 2.98; n =2) plants had ∼2-fold higher expression than their *solo* siblings (ΔCt = - 3.90; n = 2), but any genotypic differences in expression between *solo* and multi-copy plants were not significant with this small sample size (ANOVA, *P* = 0.37). Regardless, there was no evidence of 300-fold higher expression in *high+* plants.

### Copy number variants at RLG1a experience fluctuating selection across years

We examined genotype-fitness associations over three years (2015-2017) spanning the extremes of fecundity in the Iron Mountain population (seedset of plants surviving to fruitset = 29.7 ± 2.9 SE, 287.6 ± 32.2 SE, and 63.8 ± 5.5 SE, respectively; all phenotypic N > 350). In 2015, which was a record drought year in the Oregon Cascades (https://wcc.sc.egov.usda.gov) with an unusual April melt-out, we also monitored survival (< 32% overall). We recovered tissue from only ∼50% of marked seedlings (N = 243 Spring/Fall germinant pairs) and all 59 Spring germinants recovered (as dead tissue for DNA extraction) died prior to flowering. Additional flowering plants marked one week apart in mid-June also mostly died prior to setting seed (54.6% mortality, N = 185), with those that initiated flowering at the later time significantly more likely to survive (59.7% vs. 38.2%; **χ**^2^_1 df_ = 7.7, P < 0.01). RLG1a genotype was significantly associated with survival to seedset (**χ**^2^ _2 df_ = 8.56, *P* = 0.014); 64.7% (11/17) of *high+* individuals marked as seedlings or flowerers set seed, whereas only 33.0% (33/100) and 29.8% (54/181) of *trip+* and *solo* plants did. The survival advantage of *high+* plants led to significantly higher log-transformed female fitness in monitored plants (ANOVA, P = 0.005, N = 298), as they made (on average) ∼2x as many seeds (Fig. 5a). There was no significant fecundity effect in the set of random survivors collected at fruiting (P = 0.38, N = 110).

**Fig. 5.**
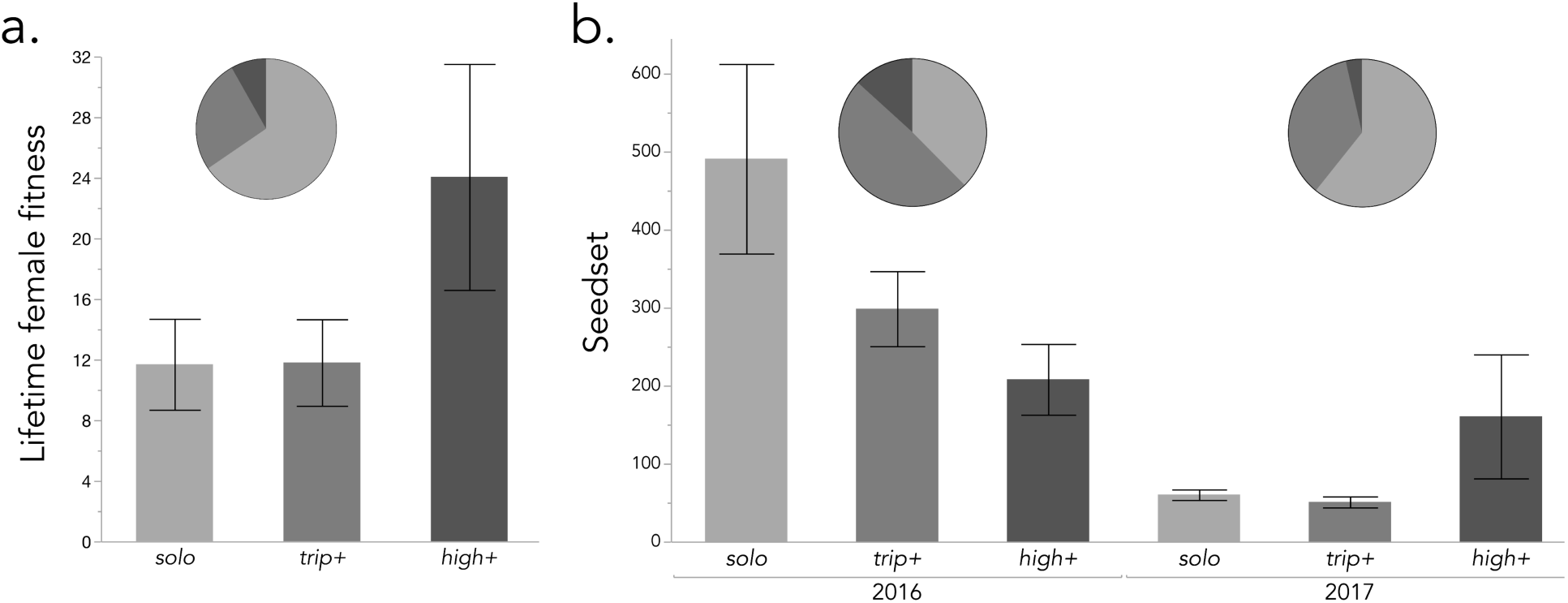
Fitness of RLG1a CNV genotypes in the field (means ± 1 SE) and frequencies of each genotype in survivors (pie charts). **a.** Lifetime female fitness (survival x seedset) in 2015, a severe drought year. **b.** Seedset (fecundity) of plants that survived to fruit in 2016 and 2017. Note different y-axis scales.

Fecundity varied significantly with RLG1a genotype in 2016 and 2017, but in opposite directions. In lush 2016, which had 5-10x the mean fecundity of flanking years, *solo* individuals matured 50% more fruits than plants with either CNV (12.4 vs. 7.7 and 8.6 for trip+ and high+ respectively), and, on average, produced twice as many seeds (Fig. 5b). In 2017, a heat wave at the end of June/early July sharply ended an otherwise lush flowering season (L. Fishman, pers. obs.). In this truncated season, *high+* plants made (on average) twice as many fruits (4.7 vs. 2.4 and 2.2) and 3 times as many seeds as the other two genotypes. These inter-annual shifts result in significant (for fruits; P = 0.016) and marginally non-significant (for seeds; *P* = 0.07) year x genotype interaction effects in the GLM, along with main effects of RLG1a genotype (both *P* < 0.005). Along with the 2015 data (and the bias toward early flowering in 2014), shifts in its relative fecundity suggest that *high+* influences life history in ways that are beneficial in stressful years (and/or microsites), but deleterious when conditions are good.

Notably, the relative proportions of the CNVs changed as predicted from the previous year’s fitness effects, consistent with real-time responses to fluctuating selection. Following disproportionate *high+* survival in 2015 (6.8% in fruiting plants), *high+* was maximally common (13%) in 2016 (when it had low seedset), and then dropped to 3.6% in 2017 (Fig. 5). Together, these results suggest that RLG1a *high+* variants have environmentally-dependent and thus temporally fluctuating effects on fitness.

## DISCUSSION

Despite the decreasing costs of sequencing, connecting phenotype to genotype remains a empirical challenge. Our original intention was to use a simple PoolSeq experiment to gain insight into genetic basis of standing variation: the focal Iron Mountain *M. guttatus* population is a place where selection should be maximally effective (and detectable) at the level of the individual nucleotide, yet also maintains tremendous diversity in fitness-relevant phenotypes. Our results provide a case study for our original question about the processes maintaining polymorphism, but also highlight the challenging complexity of both genome architecture and phenotypic expression in wild populations.

From initial Poolseq scans, we identified a tRNA ligase (RLG1a) as a coverage outlier associated with intra-population variation in flowering time, germination time, flower size, and vernalization requirement. Elevated RLG1a exon coverage in some populations and inbred lines pointed to CNV, and we identified an intermediate frequency (>14%) variant with at least three copies of the gene (*trip+).* Individual genotyping confirmed an assocation of *trip+* variants with a “slow” life-history (late flowering under stress and large flower size), and expression analyses suggest that elevated RLG1a dosage in *trip+* plants may contribute to its effects. However, even complete fixation of a 3x variant could not account for observed 10x coverage in three of our population pools, implying the existence of even higher copy-number variants. By targeting rare wild individuals carrying a marker allele sharing a diagnostic exonic indel with *trip+,* we identified a second CNV (*high+*) that carries an unprecedented 250-300 haploid copies of RLG1a in a single (presumably tandem) cluster segregating as a presence/absence polymorphism at a nuclear locus unlinked from the single-copy RLG1a. The presence of even a small number of *high+* individuals in a pool was sufficient to grossly bias initial PoolSeq scans, but individual genotyping was able to confirms significant shifts in CNV frequency. Finally, using field fitness measures from three years with very different climatic conditions, we found that *high+* experiences fluctuating selection, increasing survival and/or seed production in two drought years, but associated with low fitness relative to non-carriers in a lush year. Taken together, our discovery and analysis of the *trip+* and *high+* variants provides insight into the origins of CNV, suggests new candidate functions for an essential but poorly understood plant gene, and supports a role for climatic fluctuation in the maintenance of standing variation for fitness traits.

### An embarrassment of riches: detection and dissection of multiple layers of genic CNV

This study of just a single gene, within a single population of monkeyflowers, illustrates both the richness of copy number variation in natural populations and the significant challenges it creates for connecting genotype and phenotype (Hoban *et al.* 2016). Using genomic resequencing, qPCR, and genetic markers, we define and describe three distinct classes of RLG1a genotype, spanning at least two loci (Fig. 6). One locus, Migut.N02091 (RLG1a_1) on LG14, segregates diverse single-copy variants. A second unlinked locus (RLG1a_2) segregates for the ∼300-copy *high+* variant. The *trip+* variant(s), in which one of two inferred copes share a diagnostic 5AA deletion in exon16 with *trip+,* is likely also an allele at this locus and there are likely also duplicated (*dup*) individuals with one copy at each locus. Thus, the history of these variants includes (at minimum) duplication/ insertion of a gene into new chromosomal location, local tandem duplication and sequence divergence, and massively ampliconic tandem duplication, spanning the described mechanisms of CNV generation (Hastings *et al.* 2009). Although more work will be necessary to fully describe the evolutionary history of this extensive copy number variation, we can propose testable hypotheses about its origins.

**Fig. 6.**
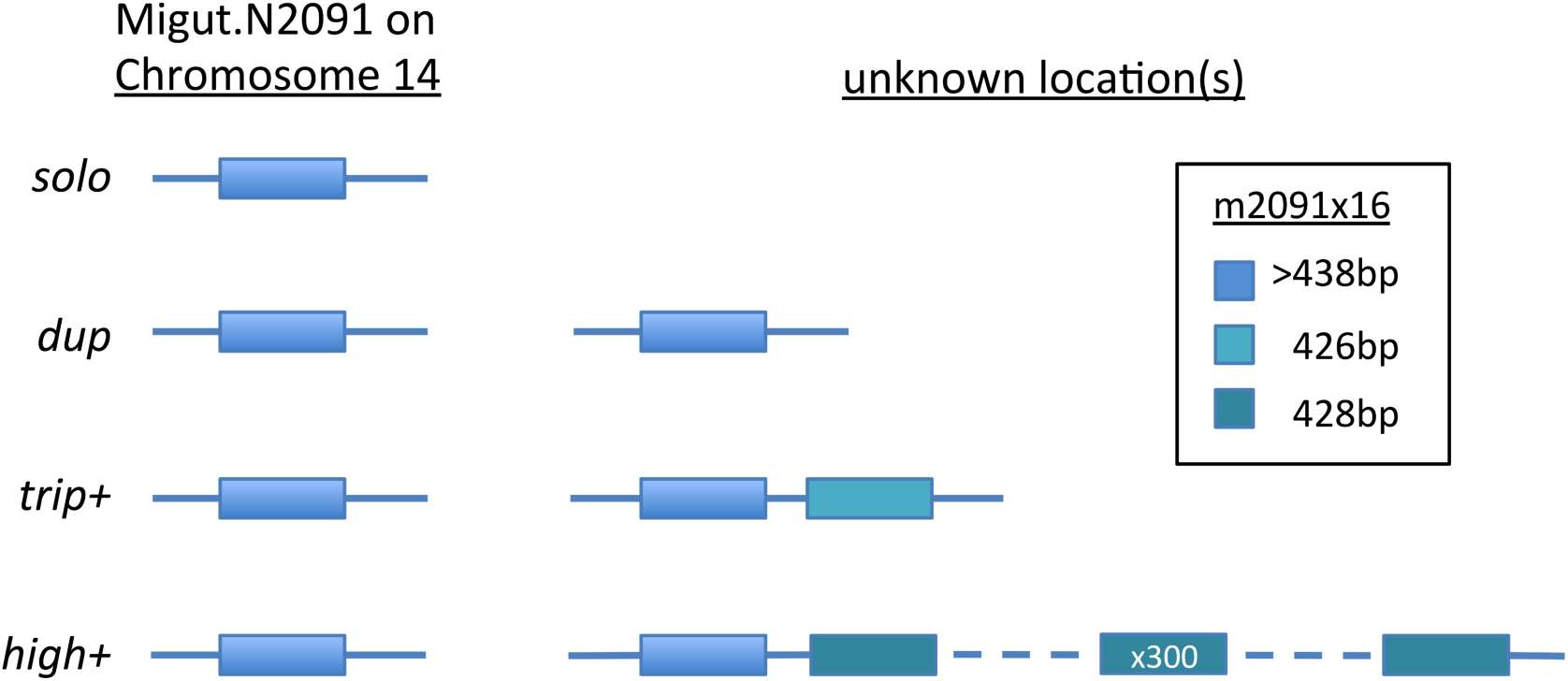
Proposed model for the structure of RLG1a copy number variation in *Mimulus guttatus*.

The tRNA ligase RLG1 is considered essential (Yang *et al.* 2017) and is the sole member of its gene family in plants (Englert & Beier 2005). Except for three Brassicaceae and *M. guttatus*, all 37 Eudicot genomes in Phytozome (ww.Phytozome.org, accessed 07/15/2018) encode just a single copy of this large gene (26 coding exons, >1200AA in *M. guttatus*). Thus, RLG1 appears highly conserved for copy number despite the multiple rounds of whole genome duplication within most Eudicot lineages. *Mimulus* appears different, even beyond the remarkable *high+* amplification. In addition to our focal locus Migut.N02091, a second tRNA ligase (Migut.D02182; RLG1b) is found on LG4. As annotated, these two genes share 82.6% amino-acid identity, suggesting a recent (most likely within *Mimulus*) duplication. Retention of two copies of RLG1 in an ancestral *Mimulus* genome may have been key to further copy number diversification within *M. guttatus*. Although the functional biology of RLG1 is still poorly understood (see next section), relaxation of the constaints imposed by a single gene performing multiple essential functions (Englert & Beier 2005) may have allowed for neo-and/or sub-functionalization (Lynch & Conery 2000) that facilitated intraspecific CNV.

The origin of copy number variants at Iron Mountain likely involved whole-gene duplication and deletion deeper in the history of the *M. guttatus* species complex. Segregation analyses show that the *high+* haplotype is present as an an unlinked locus (RLG1a2 with 250-300 *high+* copies and a putative 2 *trip+* copies; Fig. 6). This nontandem duplication is likely recent, as good agreement between estimates of copy number from qPCR amplification of a highly conserved segment and resequence read-coverage indicate high sequence identity among copies at the two loci. The diagnostic 5AA indel in exon 16 (captured by our marker mN2091×16) provides some intriguing clues as to how this may have happened. All IM individuals genotyped (including *dup, trip+* and *high+* as well as the reference and other solo genotypes) have at least one copy with the longer sequence at this position, suggesting it is ancestral. However, genome alignments from *M. nasutus* and other “Southern Clade” members of the *M. guttatus* complex (Brandvain *et al.* 2014) reveal that the deletion found in *trip+* (as one of several) and high+ (as 99% of hundreds) is the common *single-copy* variant elsewhere in the range (L. Fishman, unpublished data). Barring recurrent mutation or gene conversion, this suggests that the IM reference single copy (insertion) and the Southern clade single copy (deletion) are both derived (via reciprocal gene loss) from a duplicated ancestor similar to *trip+* IM lines (Fig. 5). Such reciprocal deletion of duplicates has recently been implicated in hybrid lethality in *M. guttatus* species complex (Zuellig & Sweigart 2018), and is a common mechanism of plant hybrid incompatibility (Fishman & Sweigart 2018), so may have implications beyond phenotypic differences of the alternative genotypes. Testing this scenario against alternatives will require genetic mapping and physical deconvolution (e.g., with long-range sequencing technologies) of *high+*, *trip+* and putative *dup* RLG1a haplotypes, as well as molecular evolution analyses of genotypes from throughout the species range.

Finally, how did the massive expansion of RLG1a gene copies in *high+* haplotypes occur? The largest genic CNV known in plants is the target gene for glyphosate herbicides, 5-enolpyruvylshikimate-3-phosphate synthase (EPSPS), which expands to 5-160 copies in *Amaranthus palmerii* (Gaines *et al.* 2010). Expansion of EPSPS confers herbicide resistance via gene dosage effects (i.e., increased expression), and is thus strongly favored in sprayed crop fields. *Amaranthus* EPSPS clusters are dispersed throughout the genome (Gaines *et al.* 2010) and cytogenetics reveals that EPSPS clusters form extra-chromosomal circular DNA (eccDNA) molecules rather than being integrated into linear chromosomes (Koo *et al.* 2018). Rolling circle amplification in eccDNA was previously implicated in the origin of the *sobo* satellite repeat in *Solanum bulbocastanum*, in which a 360kb (nongenic) monomer amplified to 4.7 Mb in a single (hemizygous) chromosomal location (Tek *et al.* 2005). RLG1a *high+* segregates as the presence/absence of a tandem array, and thus may have originated via rapid amplification in eccDNA and re-insertion (without copy number intermediates) into the nuclear genome (Charlesworth *et al.* 1994). More work is necessary to evaluate this scenario, and to test the effects of RLG1a2 expansion on the recombination and segregation of linked loci.

This study highlights how much CNVs complicate genome scans for adaptive differentiation, despite being a large proportion of potentially functional variants. Especially in pooled population sequencing (PoolSeq) experiments, CNV generates substantial complications. For example, Migut.N02091 outlier sites were censored (for excess coverage) in SNP scans of the High-Low flower size selection experiment (Kelly *et al.* 2013), but it appeared to be strong SNP outlier in the IM flowering time study (Monnahan & Kelly 2017). Sites with low coverage are rightly dropped from most analytical pipelines (DePristo *et al.* 2011), so gene deletions relative to a reference genome (particularly within clusters of paralogs where cross-mapping may occur) are difficult to identify and such loci may be lost from scans. Particularly in PoolSeq studies, the coverage stacks created by duplication can generate false positives via statistical (over-confidence) and biological bias (mis-representation of allele frequencies). Summary statistics of differentiation between pooled populations reflect sample size (coverage) as well as allele frequencies (Kofler *et al.* 2011; Magwene *et al.* 2011; Kelly *et al.* 2013), so even identical duplicates can inflate statistical significance. At the same time, stacking of multiple paralogs with shared SNP/indel differentiation (like *trip+* and *high+ vs. solo)* exaggerates differences among pools. Without the 2bp indel that happens to distinguish *trip+* and *high+* diagnostic PCR amplicons (and follow-up targeted resequencing, crossing, and qPCR), we would have parsmoniously inferred the existence of a 5-copy variant much more differentiated in frequency among pools. Distortion of PoolSeq SNP scans by CNV adds to the existing arguments for using large sample sizes (>100 individuals) in PoolSeq experiments (Lynch *et al.* 2014). In addition, validation of results from genome scans (using both individual and pooled sequencing approaches) with independent genotyping is an important step in investigating the genetic basis of adaptive traits.

Along with the increasing evidence that CNV contributes to fitness-relevant variation in crops and humans (Girirajan *et al.* 2011; Maron *et al.* 2013; Mickelbart *et al.* 2015; Dolatabadian *et al.* 2017), our results suggest that explicit CNV surveys should be routine part of genome-wide scans for adaptive loci. A number of new pipelines combine coverage, read-pair information, and other data to infer structural variants from short-read data (Layer *et al.* 2014; Mohiyuddin *et al.* 2015), and could fruitfully be used to catalog CNV in (for example) the Iron Mountain inbred line set. The detection of putative 2x-covered *dup* individuals in that set (Fig. 2) is promising, though poor alignment in intergenic regions in diverse *M. guttatus* limits the utility of read-pair information as corroborative evidence. Without such a reference catalog of variants (or until the advent of inexpensive single-molecule sequencing applicable at a population scale), however, whole-gene CNVs are likely to often remain a component of the missing heritability (Manolio *et al.* 2009).

### One piece of the puzzle: a tRNA ligase and standing variation for life-history traits

The genic CNVs identified in this study are undoubtedly a tiny fraction of the variants affecting life-history phenotypes in IM *M. guttatus*, but RLG1a is the first (and hopefully worst!) gene to be dissected mechanistically as a direct contributor to fitness phenotypes in this model population. Importantly, despite not knowing the chromosomal locations of the CNVs or full individual genotypes, we can be reasonably confident that RLG1a copy number *per se* and/or sequence variation within the characterized CNVs underlies the observed phenotypic and fitness associations. In IM *M. guttatus*, linkage disequilibrium (LD) falls off sharply at distances >1000bp (with the exception of several inversions and/or recent selective sweeps; Puzey *et al.* 2017). Thus, *trip+* and *high+* are unlikely to be neutral markers for functionally-distinct loci tightly linked to RLG1a2; in addition, a preliminary scan for associated sites (i.e, scanning for differences in nucleotide diversity within vs. between *solo* and *trip+* inbred line sets; data not shown) did not reveal any strong or extensive LD with other genomic regions. Thus, we discuss the phenotypic and fitness results in the context of RLG1 function.

Why a tRNA ligase CNV? Plant tRNA ligases share little sequence homology with tRNA ligases in other taxa, but retain the core function of splicing the two intron-containing nuclear-encoded tRNAs (tRNA^Tyr^ and elongator tRNA^Met^) (Englert & Beier 2005). The single-copy *Arabidopsis* RLG1 is expressed in most tissues and is embryo-lethal when knocked out (Yang *et al.* 2017). RLG1 has also been implicated in the joint regulation of the key plant hormone auxin and the endoplasmic reticulum-based unfolded protein response (UPR) in *Arabidopsis* (Leitner *et al.* 2015; Nagashima *et al.* 2016). One arm of the UPR is regulated by cytoplasmic splicing of bZIP60 transcription factor mRNAs; RLG1 performs the key ligation step (Nagashima *et al.* 2016). Via its regulation of the UPR, RLG1 may play a central role in plant reactivity and resistance to abiotic stressors such as heat and drought, as well as hormonal control of tradeoffs between vegetative growth and reproduction (Deng *et al.* 2013). In addition, the bZIP60 branch of the UPR is constitutively up-regulated in pollen (which is notoriously vulnerable to heat stress) (Deng *et al.* 2016) and integral to plant Reponses to viral infection (Zhang *et al.* 2015). Even beyond its remarkable copy number diversity, *M. guttatus* (with both RLG1a and RLG1b in everyone) provides a rare opportunity to investigate the multiple functions of plant tRNA ligases without the constraint of an essential single copy.

Our phenotype and fitness results suggest that RLG1a CNVs contribute to *stress-responsive* variation in the timing of reproduction, along a spectrum from highly reactive to stress (fast) to relatively nonchalant (slow). Interestingly, carriers of the *trip+* variant appear “slow” (large flowers, later reproduction in field and under drought stress), whereas *high+* carriers appear “fast” (early flowering in 2014, highest survival and/or seedset in drought years). Further analyses of sequence variants and expression patterns will be necessary to determine the physiological mechanisms underlying this pattern. Importantly, the fecundity effects of RLG1a CNV that we see in the field are caused by differences in fruit number rather than seedset/fruit (which did not significantly differ among years or RLG1a genotypes). Thus, fluctuating fitness effects depend on life-history plasticity/stress tolerance traits that affect the timing of flowering and senescence. This parallels earlier work showing fluctuating selection on “fast” and “slow” QTL introgression lines derived from the flower size selection experiment (Mojica & Kelly 2010; Mojica *et al.* 2012), but highlights genotype x environment interactions for phenotype as well as fitness. For example, alternative RLG1a genotypes may differ in whether they bolt faster when sensing short-term heat stress via the UPR, with the fitness consequences depending on whether heat stress accurately predicts imminent dry-down and death in a given year and microsite. In addition to mediating stress-responsive cues, RLG1a CNVs could also affect stress-tolerance; elevated expression of RLG1a (as in *trip+* lines; Figure 3) may increase the efficiency of UPR-upregulation and thus protect against heat or cold damage that shortens life span. Unmeasured effects on male fitness (e.g., via heat tolerance of pollen) may also contribute to the maintenance of alternative RLG1a CNVs. Exploring these intriguing possibilities in the appropriate environmental contexts will require long-term monitoring of natural populations, as well as experimental manipulation of candidate stress regimes.

Temporally fluctuating selection is an appealing but controversial mechanism to maintain fitness variation, as simple models cannot indefinitely maintain multiple alleles when their fitness effects are not perfectly balanced (reviewed in Delph & Kelly 2013). However, plasticity (which we observe) and shifts in dominance (which are implied by *high+/solo* fitness reversals; Fig. 4) can enlarge the parameter space over which temporal variation prevents allele loss (Posavi *et al.* 2014; Gulisija *et al.* 2016; Wittmann *et al.* 2017). In addition, dormant life-stages (e.g. a seedbank) and spatial variation within a population (e.g., unpredictable soil depth) can slow the dynamics of displacement and thus help maintain standing variation over observable time scales (Ellner & Hairston 1994; Svardal *et al.* 2015). Although we cannot be sure that *high+* will not either fix or be driven from the Iron Mountain population in the long term (especially as it may be a relatively new variant not at equilibrium), the observed shifts in relative fitness from year to year plausibly contribute to its maintenance at low to intermediate frequency. This parallels the classic *Linathus parryae* system, in which temporal and spatial variation in drought stress contributes to the maintenance of a flower color polymorphism (Schemske & Bierzychudek 2001; 2007). In a model inspired by that system, variants with geometric mean fitness < 1 (i.e., ones that should be deterministically lost otherwise) can be maintained if they have highest fitness in “good” years and a seedbank spreads that advantage over time (Turelli *et al.* 2001). RLG1a in the IM *M. guttatus* population plausibly experiences similar dynamics, mediated (in part) by stress-responsive phenological traits. Our results add to the growing evidence that temporally fluctuating selection cannot be empirically irrelevant to the abundant quantitative genetic variation seen in many short-lived taxa (Siepielski *et al.* 2009).

## ACKNOWLEDGMENTS

We thank Gloria Goni-McAteer (2015), Hanna McIntosh and John Crandall (2016), and Emily Beck (2017) for helping with the field plant collections, and Daniel Crowser, Findley Finseth, and Katie Zarn for lab and greenhouse assistance. We appreciate the logistical support for library preparation, qPCR, and marker genotyping provided by Tamara Max and Denghui (David) Xing in the University of Montana Genomics Core. This work was supported by NSF DEB-0846089, DEB-1457763, and OIA-1736249 to L.F. and by NIH R01-GM073990 to J.K.K.

## DATA ACCESSIBILITY

- DNA sequences (fastqs): NCBI SRA: Inbred lines: SAMN05852485-SAMN05852522, SAMN04517335; Iron Mountain flowering time pools: PRJNA336318. Fastqs for other pools and wild accessions will be archived prior to publication.
- Genotype and phenotype data, as well as qPCR datsets, will be archived on Dryad prior to publication

## AUTHOR STATEMENT

L.F. and J.K.K designed experiments, P.M., L.F. and J.K.K collected PoolSeq samples, P.M, J.K.K, F.R.F., and T.N. constructed genomic libraries and analyzed sequence data, L.F., F.R. F., T.N, and K.A. conducted qPCR experiments, L.F., M.M, and E. M-W. collected phenotypic data and PCR-genotyped wild and greenhouse-grown plants. L. F. wrote paper with input from all authors.

